# Dual role of Pectin Methyl Esterase activity in the regulation of plant cell wall biophysical properties

**DOI:** 10.1101/2022.06.14.495617

**Authors:** Marçal Gallemí, Juan Carlos Montesinos, Nikola Zarevski, Jan Pribyl, Petr Skládal, Edouard Hannezo, Eva Benková

## Abstract

Acid-growth theory has been postulated in the 70s to explain the rapid elongation of cells in response to plant hormone auxin. More recently, it has been demonstrated that activation of the proton ATPs pump (H^+^-ATPs) promoting acidification of the apoplast is the principal mechanism through which hormones like auxin as well as brassinosteroids (BR) induce cell elongation. However, the impact of this acidification on the mechanical properties of the cell wall remained largely unexplored. Here, we use Atomic Force Microscopy (AFM) to demonstrate that acidification of apoplast is necessary and sufficient to induce cell elongation through cell wall relaxation. Moreover, we demonstrate that Pectin Methyl Esterase (PME) can induce both cell wall softening or stiffening in extracellular calcium dependent-manner and that tight control of PME activity is required for hypocotyl elongation.

## INTRODUCTION

Plants, as sessile organisms, developed exceptional plasticity of growth and development to adapt to their ever-changing environment. One of the most prominent mechanisms to ensure adjustability is a rapid elongation growth that allows plant organs to reach a position optimal for utilization of light, nutrients and survival. Plant cells grow by anisotropic expansion, which results from the interplay between turgor pressure and a fine-tuned local cell wall (CW) relaxation (Wolf et al., 2012; Braidwood et al., 2014).

Plant hormone auxin is a principal endogenous regulator of elongation growth, which has been shown to induce cell expansion in plant organs such as stems, coleoptiles, or hypocotyls segments within minutes (Rayle and Cleland, 1992; Fendrych et al., 2016). The current model of auxin-regulated cell expansion is based on the acid growth theory (reviewed in Mockaitis and Estelle, 2008; Perrot-Rechenmann, 2010). In this model, auxin induces transcription of *SAUR* (*Small Auxin Upregulated genes*) genes, which indirectly activate the proton-pumping plasma membrane (PM)-H^+^-ATPase, thereby enhancing the transport of protons into the extracellular space. Recently, it has been shown that auxin can regulate PM-H^+^-ATPase through yet another mechanism mediated by Receptor-like Transmembrane Kinase1 (TMK1) (Lin et al., 2021). At low apoplastic pH, the activity of CW remodeling proteins such as expansins is enhanced, which leads to the CW relaxation (Cosgrove, 2018). In parallel to CW acidification, auxin in the nuclei enhances the transcription of genes including those encoding PM-ATPase, K^+^ channels, expansins, and other CW remodeling enzymes (Majda and Robert, 2018; Velasquez et al., 2016). Similarly to auxin, another class of plants hormones, brassinosteroids, has been reported to induce hypocotyl elongation through activation of PM-H^+^-ATPase and regulating the expression of several CW remodeling enzymes (Caesar et al., 2011; Minami et al., 2019).

Pectins are components of the CW considered to be principal determinants of its bio-physical properties (Saffer, 2018; Shin et al., 2021). They are polymers composed of α-D-galactosyluronic acid (Gal-A) forming homogalacturonan (HG) linear chains. Pectins are by default transported in a methyl-esterified state to the apoplast, where they can be de-esterified by Pectin Methyl-Esterases (PME) proteins, in a process that liberates protons. The activity of PMEs is fine-tuned by pectin methyl-esterase inhibitors (PMEIs). *In vitro* assays showed that at low pH interaction of PMEIs with PMEs is stabilized and the PME activity attenuated (Hocq et al., 2017a). De-methylesterified pectins have free carboxyl groups that are hypothesized to either form cross-links mediated by calcium ions (the so called egg-box hypothesis), which might increase the rigidity of the CW (Hocq et al., 2017b), or undergo cleavage to shorter HGs chains resulting in the softening of the CW (Cosgrove, 2022). Based on super-resolution microscopy, it has been described that pectins possibly form nanofilaments that alter their quaternary structure upon de-methylesterification and calcium cross-linking, supporting the egg-box model (Haas et al., 2020).

Despite the importance PMEs and PMEIs in the modulation of the pectin structure, there is still lack of mechanistic understanding of their functions in the regulation of cell growth processes and contradictory results have been published on the effects of those proteins (Jonsson et al., 2022; Cosgrove, 2022). Whilst some studies have reported increased CW stiffness in epidermal cells of hypocotyls overexpressing *PME5* (Bou Daher et al., 2018), others have shown that enhanced PME5 activity results in decreased stiffness of the CW (Peaucelle et al., 2011, 2015). Lines overexpressing *PMEIs* seem to raise controversy as well. Overexpression of *AtPMEI3* has been associated with increased stiffness of hypocotyl epidermal CWs, but also softening effects on CWs have been reported, both correlating with reduced cell length and growth anisotropy (Bou Daher et al., 2018; Peaucelle et al., 2008, 2011, 2015). Arabidopsis plants with increased levels of *AtPMEI2* show enhanced growth and longer root as a result of promoted cell elongation (Lionetti et al., 2007, 2010; Liu et al., 2018). Expression of *AtPME1* in tobacco pollen tubes inhibited elongation, whereas *AtPMEI2* expression led to an increase in their elongation rate (Röckel et al., 2008). Thus, it seems that a developmental or cellular context needs to be taken into account to explain the role of PME and PMEI in growth processes, their impact on CW mechanical properties and cell elongation.

Atomic Force Microscopy (AFM) techniques have recently aroused as an important tool to monitor bio-physical properties of plant CW. Here, we applied AFM to investigate how selected plant hormones, known for their principal role in the regulation of cell elongation, affect CW mechanical properties. We further applied this technique to address a question about how modulation in PME activity affects CW mechanics in relation to calcium levels in the apoplast. We demonstrate a dual role of PME activity, indicating that a narrow window of pectin methylesterification is important for proper cell elongation.

## RESULTS AND DISSCUSION

### Apoplastic acidification is necessary and sufficient to induce cell elongation through CW relaxation

Modulation of hypocotyl elongation growth is one of the prominent adaptive responses to changes in environmental conditions such as temperature or light intensity. At the molecular level, hypocotyl growth is governed by auxin, which in concert with other hormonal pathways including brassinosteroids (BR) or gibberellins (GA), promotes cell expansion. Acidification of the apoplast triggered by auxin is one of the essential regulatory steps in this process (Oh et al., 2014). While the promoting effect of acidification on hypocotyl expansion is firmly established, its impact on CW bio-physical properties is still poorly understood. Old reports using extensometers had shown that stems are able to stretch more after apoplastic acidification (Cosgrove, 1993). However, the latest results point out that tensile stretching might not correlate with CW mechanical properties quantified by other techniques like Apparent Young Modulus (*E_a_*) obtained from AFM (Zhang et al., 2019).

To establish experimental conditions to investigate the impact of apoplastic acidification on cell elongation and CW bio-physical properties, we adopted an assay using hypocotyl segments of etiolated Arabidopsis seedlings (Schenck et al., 2010; Fendrych et al., 2016). Initially, employing the ApopH apoplastic pH sensor (Gjetting et al., 2012), we confirmed that incubation of etiolated hypocotyl segments of 3-day-old seedlings in either acidic (pH 3.5) or alkaline (pH 9.0) buffers for 2 hours (h) resulted in expected changes of the apoplast pH in hypocotyl epidermis (Fig. S1A, B). Comparably to the incubation in acidic buffer, treatment with 10 μM synthetic auxin 1-Naphthylacetic acid (NAA) at pH 6.0 for 2 h led to acidification of the apoplast (Fendrych et al., 2016), while auxin was not able to trigger acidification of apoplast when the buffer of pH 9.0 was used (Fig. S1A, B).

Measurements of hypocotyl length showed that 3 h of incubation in the acidic buffer was sufficient to trigger elongation in a similar range as hormone auxin or BR (epibrassinolide, eBL) at pH 6.0. Alkaline buffer did not promote hypocotyl elongation and interfered with hormone-induced elongation (Fig. 1A, B). Importantly, analysis of the auxin-sensitive reporter R2D2 (Liao et al., 2015) showed that auxin response is not affected by the alkaline pH itself (Fig. S1 C,D).

**Figure 1.**
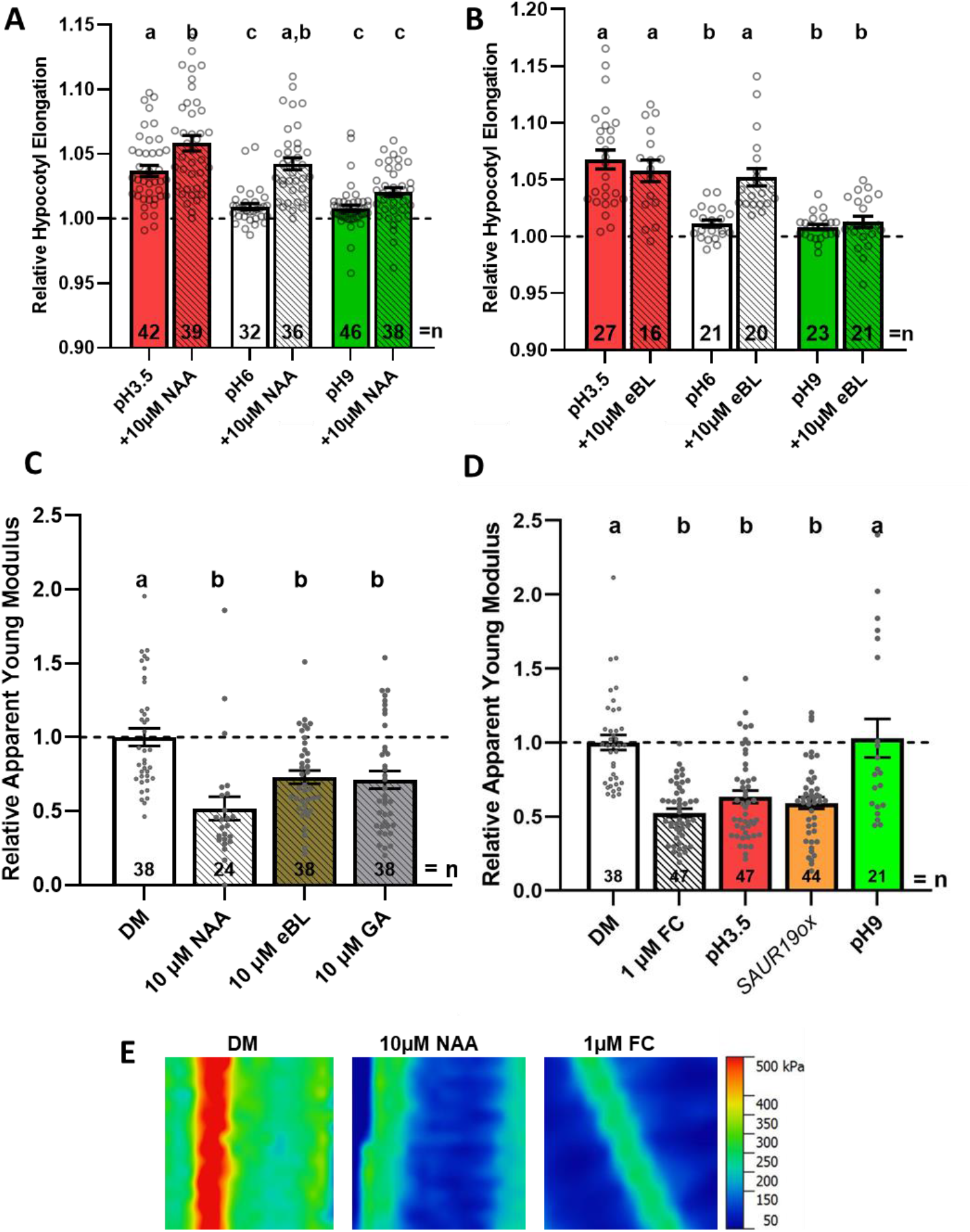
Acidification induces hypocotyl segments elongation and CW softening. A, B. Relative hypocotyl segment length after 3 h of indicated treatments. Hypocotyl segments of 3-day-old seedlings were incubated in depletion medium (DM) adjusted to pH 3.5, 6.0 or 9.0 with or without auxin (10 μM 1-naphthaleneacetic acid, NAA, panel A) and BR (10 μM epibrassinolide, eBL, panel B). Dashed lines indicate relative initial length of hypocotyl segments. C-E. Relative Apparent Young Modulus (*E_a_*) (C,D) and corresponding representative heat-maps (E) of longitudinal epidermal CWs in hypocotyl segments of 3 day-old seedlings incubated for 2 h in depletion medium (DM) adjusted to pH 6.0 supplemented with auxin (10 μM NAA), brassinosteroid (10 μM eBL), gibberellin (10 μM GA) (C) and in DM adjusted to pH 3.5, pH 9.0, supplemented with fusicoccin (1 μM FC in DM, pH 6.0), and SAUR19ox (in DM, pH 6.0) (D). Dashed lines indicate normalized *E_a_* respect to DM. Bars represent average ±*SE*, dots represents individual data points. N indicates the total number of hypocotyls segments quantified (A,B) and the total number of scans performed and analyzed (C,D). Tukey-Kramer test was performed and significant differences are shown as the letters above each bar.

Thus, consistently with previous findings, our experimental set-up using Arabidopsis hypocotyls shows that the acidification of the apoplast, achieved either through incubation in acidic buffer or hormonal treatments, is both sufficient and necessary for hypocotyl cell elongation (Cleland et al., 1991; Rayle and Cleland, 1992; Arsuffi and Braybrook, 2018).

To establish and validate our AFM experimental platform for measurements of CW mechanical properties, we inspected how CW rigidity of hypocotyl epidermal cells is altered in different growth regimes. Hence, we measured *E_a_* in either long etiolated, or short de-etiolated hypocotyls of wild-type (Col-0) or hypocotyls of light insensitive mutants in phytochrome A and B (*phyAphyB*), which elongate despite growing in light conditions (Fig. S2A). We observed a strong correlation between CW mechanical properties and hypocotyl elongation. The average apparent *E_a_* measured on longitudinal CWs (parallel with hypocotyl growth axis) in etiolated hypocotyls was around 441 ±87 kPa *Standard Error* (*SE*). In light-grown hypocotyls *E_a_* increased to 3699 ±670 kPa, while in the *phyAphyB* mutant (with intermediate hypocotyl elongation) intermediate *E_a_* values of around 1544 ±130 kPa were detected (Fig. S2B, C). Those results are largely in agreement with previously published reports (Peaucelle et al., 2015), including measurements using other techniques such as Brillouin scattering microscopy (Elsayad et al., 2016).

Applying the same AFM set-up, we analyzed the impact of hormonal treatments on the mechanical properties of CWs in hypocotyl segments. Measurements of cell growth confirmed that epidermal cells in the middle part of hypocotyl segments of 3-day-old seedlings rapidly elongate in response to auxin when compared to untreated control (Fig. S3). Thus, analyses by AFM were standardly performed on epidermal cells in the middle zone of hypocotyl segments.

All treatments, which enhanced hypocotyl elongation including auxin, BR and GA, decreased CW stiffness at a similar range (60% of *E_a_* compared to the non-treated hypocotyls; 0.52 ±0.08 *SE* for auxin, 0.72 ±0.05 for BR and 0.71 ±0,06 for GA; Fig. 1C-E). As hormones such as auxin and BR can induce apoplastic acidification (Fendrych et al., 2016; Minami et al., 2019), we inspected whether direct acidification of apoplast would induce similar changes in CW stiffness. Apoplastic acidification as a result of incubation of hypocotyl segments in acidic buffer (pH 3.5) for 2 h, or the PM-H^+^-ATPase pump activation by either fusicoccin drug (FC, Baunsgaard et al., 1998; Ballio et al., 1964) or overexpression of *SAUR19ox* (Spartz et al., 2012) led to the significant reduction of *E_a_* when compared to wild-type control incubated in the buffer of pH 6.0 (0.67 ±0.06 for pH 3.5 buffer, 0.53 ±0.03 for FC, and 0.59 ±0.04 for *SAUR19ox*). The incubation in an alkaline buffer had no significant effect on the *E_a_* of CW in hypocotyl epidermal cells (Fig. 1D, E). Hence, we hypothesize that various hormonal treatments, which induce hypocotyl elongation, converge on the regulation of mechanisms that control acidification of the apoplast and subsequently promote CW softening.

### Auxin-induced softening of the CW requires intact signaling pathway

Our results indicated that hypocotyl elongation triggered by hormones like auxin or BR or apoplast acidification correlates with CW softening. Published works have shown that an intact signaling pathway is required for auxin-induced acidification and hypocotyl elongation (Fendrych et al., 2016), but that BR is able to activate PM-H^+^-ATPs directly (Caesar et al., 2011; Minami et al., 2019). Using cycloheximide (CHX), an inhibitor of proteosynthesis (Rose, 1974), in agreement with previous studies, we observed that auxin-mediated hypocotyl elongation requires protein biosynthesis (Bates and Cleland, 1979; Rayle and Cleland, 1980; Kutschera and Schopfer, 1985; Fendrych et al., 2016), whereas hypocotyl elongation triggered by incubation in acidic buffer (pH 3.5) is not dependent on *de-novo* protein synthesis (Fig. 2A,B). Interestingly, BR promoting effect on hypocotyl elongation is only partially independent of *de novo* protein synthesis. Hypocotyls elongate more if proteosynthesis is not compromised, suggesting that in addition to direct activation of the proton pump another protein biosynthesis dependent mechanism might contribute to the regulation of hypocotyl elongation (Fig. 2A,B). Next, we examined how inhibition of auxin signaling pathway affects CW mechanical properties and sensitivity to auxin. Using a heat-inducible *HS::axr3-1* transgenic line (Knox et al., 2003), we found that accumulation of *axr3-1*, a dominant negative repressor of auxin signaling and auxin-induced hypocotyl elongation (Fendrych et al., 2016), interferes with auxin-mediated softening of CWs when compared to no-heat treated control (Fig. 2C). We conclude that auxin-induced hypocotyl elongation and CW softening are dependent on new protein biosynthesis and intact transduction cascade.

**Figure 2.**
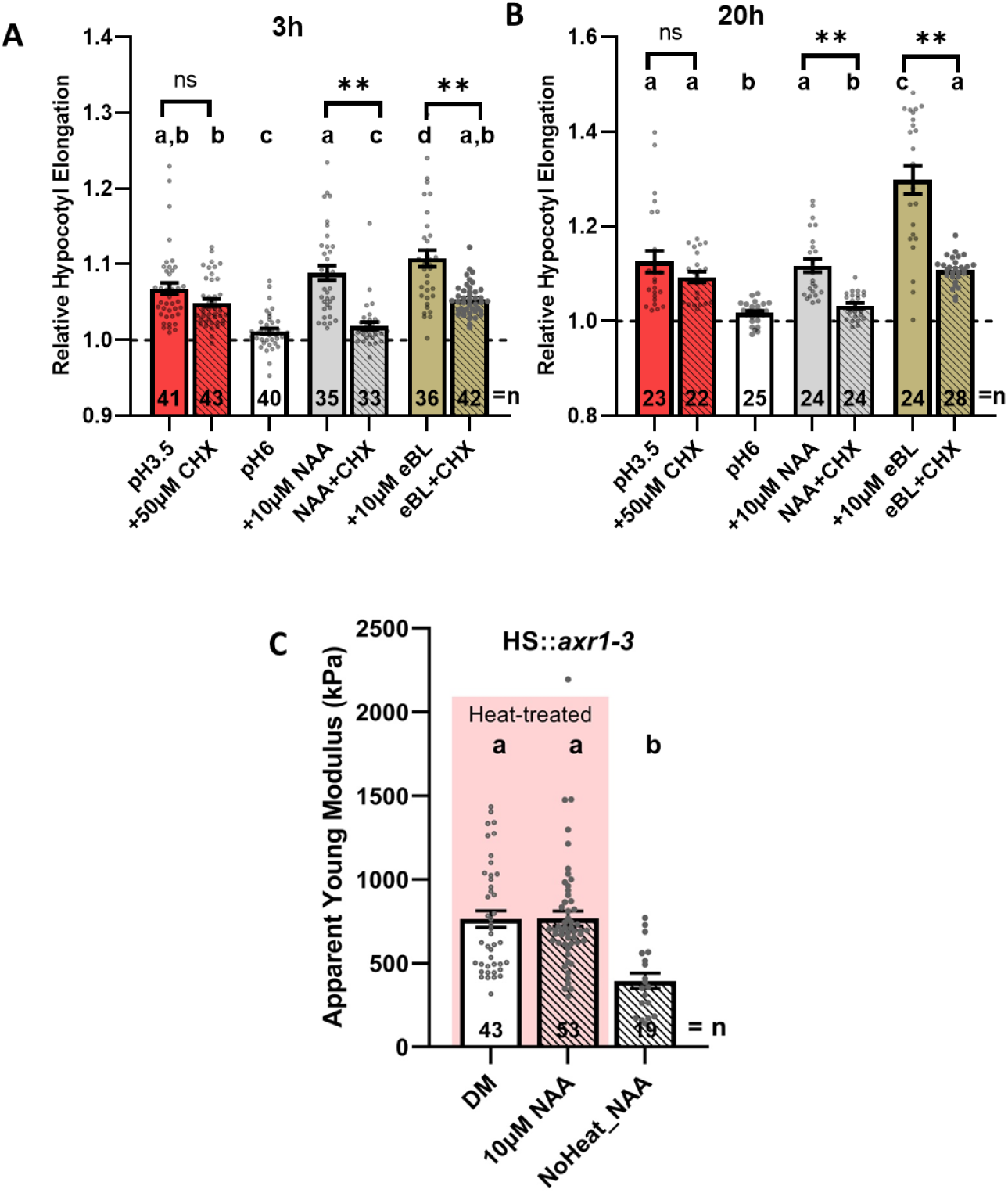
Auxin-triggered hypocotyl elongation is proteosynthesis dependent. A, B. Relative hypocotyl segment length after 3 h (A) or 20 h (B) of indicated treatments. Hypocotyl segments of 3 day-old seedlings were incubated in DM medium adjusted to pH 6.0 or 3.5 (red bars) and supplemented with hormones (NAA and eBL) with or without the protein synthesis inhibitor cycloheximide (50 μM CHX), respectively, as indicated. Dashed lines mark initial relative length of hypocotyl segments. C. Average *E_a_* quantification of longitudinal epidermal CWs of 3 day-old hypocotyl segments of *HS::axr3-1* seedlings, non- or heat treated for 40 min at 37 °C, respectively, and incubated in DM with or without auxin for 2 h. Bars represent average ±*SE*, dots represents individual data points. N indicates the total number of hypocotyls segments quantified (A and B) or scans performed and analyzed (C). Tukey-Kramer test was performed and significant differences are shown as the letters above each bar. Direct pair significant differences are indicated as **P < 0.01 (t-test).

### Balanced PME activity mediates fast hypocotyl elongation

We showed that hormone-induced apoplastic acidification is necessary for fast hypocotyl elongation and CW softening. However, processes associated with the re-structuring of CW that result in modulation of its biophysical properties are still poorly understood. Pectins, as important components of the primary CW, have been proposed as principal determiners of the mechanical properties of the CW (Shin et al., 2021; Saffer, 2018). In particular, a degree and pattern of methylesterification of homogalacturonan chains controlled by Pectin Methyl Esterases (PME) and PME Inhibitors (PMEI) antagonizing their activities, might have a decisive impact on CW mechanics (Hocq et al., 2017b).

To study early responses to transient increase of PME activity, we generated inducible *PME* overexpressing (*ox*) lines. *PME1* as an example of the auxin-inducible homolog of the PME family (Vanneste et al., 2005; Nemhauser et al., 2006; Simonini et al., 2017) and a previously studied *PME5* were selected for detailed analyses (Peaucelle et al., 2015; Bou Daher et al., 2018). Several independent lines were obtained and selected based on expression levels determined by RT-qPCR and Western-Blot (Fig. S4A and B). The expression of *PME1ox* as well as *PME5ox* in seedlings exposed to β-estradiol (β-EST) for 48 h affected seedling development and resulted in shorter roots compared to non-induced seedlings (Fig. S4C), indicating that recombinant PME1-HA and PME5-HA proteins maintain their activities.

To examine how enhanced expression of *PMEs* affects CW mechanical properties we employed AFM. Monitoring of CW stiffness in hypocotyl epidermal cells revealed surprising, time-dependent effects of PMEs on the CW mechanics. Induction of *PMEs* expression for a short time (~24 h) resulted in stiffer CWs, whilst *PMEs* expression persisting for about 3 days correlated with CW softening when compared to untreated control (Fig. 3A). Enhanced expression of *PME5* has been reported to result in stiffening of CWs in hypocotyl epidermal cells (Bou Daher et al., 2018), while in other study softening of hypocotyl epidermal cells and shoot apical meristem has been detected (Peaucelle et al., 2011, 2015). In light of our observations, we hypothesize that the previously observed softening of CWs might be a result of prolonged and/or very strong PME activity. As specified by the authors (Peaucelle et al., 2015, supplemental information) seedlings with high expression and showing strong phenotype defects were selected to analyze. In our experiments, we did not apply any pre-selection based on phenotypes.

**Figure 3.**
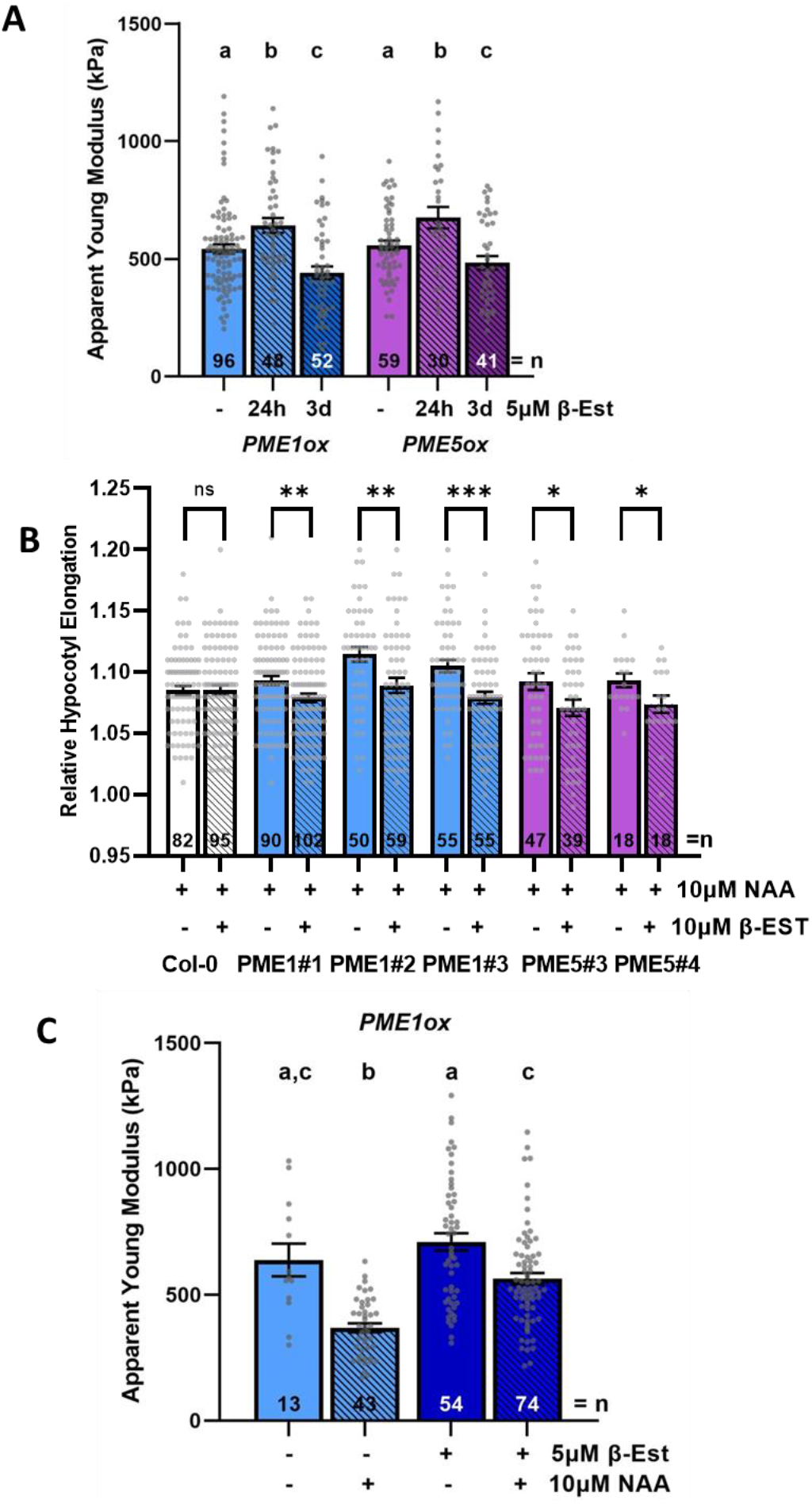
Enhanced expression of *Pectin Methyl-Esterase1* (*PME1*) or *PME5* modulates CW stiffness and interferes with auxin promoted hypocotyl elongation. A. Average *E_a_* quantification of longitudinal CWs measured in hypocotyl epidermal cells of 3 day-old seedlings of *PMEoxs* lines at indicated induction time with β-estradiol (β-Est). B. Relative hypocotyl segment length in wild-type Col-0, and *PME1ox* and *PME5ox* lines non- or treated with β-Est for 24 h and auxin (1-naphthaleneacetic acid, NAA) for 3 h, as indicated. C. Average *E_a_* quantification of longitudinal CWs of hypocotyl epidermis in 3 day-old *PME1ox* seedlings non- or treated with β-EST for 24 h and hypocotyl segments non- or treated with auxin (NAA) for 2 h. Bars represent average ±*SE*, dots represents individual data points. N indicates the total number of hypocotyls segments quantified (B) or scans performed and analysed (A and C). Tukey-Kramer test was performed and significant differences are shown as the letters above each bar. Direct pair significant differences are indicated as *P < 0.05, **P < 0.01, and ***P < 0.001 (t-test).

Next, we analyzed how the enhanced expression of *PME1* or *PME5* affects auxin-induced elongation. Intriguingly, growth response to auxin was hampered in hypocotyls in which *PMEs* expression was induced (24 h of induction by β-EST and 3 h treatment with auxin) when compared to non-induced controls treated with auxin only, indicating that modulation of pectin methylesterification by PMEs interferes with auxin-triggered elongation of hypocotyls (Fig. 3B). As an enhanced *PME* expression interfered with auxin-induced growth, we tested whether this PME effect correlates with alteration of CW mechanical properties. AFM measurements revealed that *PME* expression significantly attenuates auxin capacity to relax CWs (Fig. 3C).

Altogether, these results suggest that tightly controlled pectin de-methylesterification is required for fast hypocotyl elongation in response to auxin, and that modulation of the PME expression might lead to either softening or stiffening of CW, depending on the duration of the PME activity.

### Calcium availability affects the PME mediated CW stiffening

Recently, the paradox of how PME activity might lead to two distinct effects on CW mechanics has been discussed and a model proposed, in which a pattern of pectin de-methylesterification is an important determiner of CW stiffness (Hocq et al., 2017a, 2017b; Arsuffi and Braybrook, 2018). Blockwise HGs demethylation might lead to the formation of stretches of negatively charged HG chains that can interact with divalent Ca^2+^ ions and form the so-called egg-box structures, increasing the stiffness of CWs. On other hand, random de-methylesterification might enhance cleavage of HGs chains by pectin-degrading enzymes to oligogaracturonides and thereby promote CW relaxation.

Our AFM measurements in hypocotyls pointed out that CW biophysical properties can be modified differently in dependence on the duration of PME activity (Fig. 3A). As Ca^2+^ ions appear to be a critical factor determining the rigidity of CWs, we examined whether modulation of Ca^2+^ levels affects CW mechanical properties and hypocotyl capacity to elongate. To explore whether increased CW stiffness as a result of enhanced *PME1* expression might involve Ca^2+^ mediated cross-linking of HGs, we incubated hypocotyls of *PME1ox* line in the medium supplemented with calcium chelator ethyleneglycoltetraacetic acid, (EGTA, Feng et al., 2018). No increase in CW stiffness was detected in hypocotyl epidermal cells of *PME1ox* induced by β-EST incubated in the medium supplemented with EGTA when compared to non-EGTA treated controls. The CW stiffness in hypocotyls overexpressing *PME1ox* and treated with EGTA was comparable to that detected in control non-induced hypocotyls (Fig. 4A). These results suggest that the availability of Ca^2+^ might be critical for cross-linking of de-methylesterified HGs shortly after activity of *PME* is increased.

**Figure 4.**
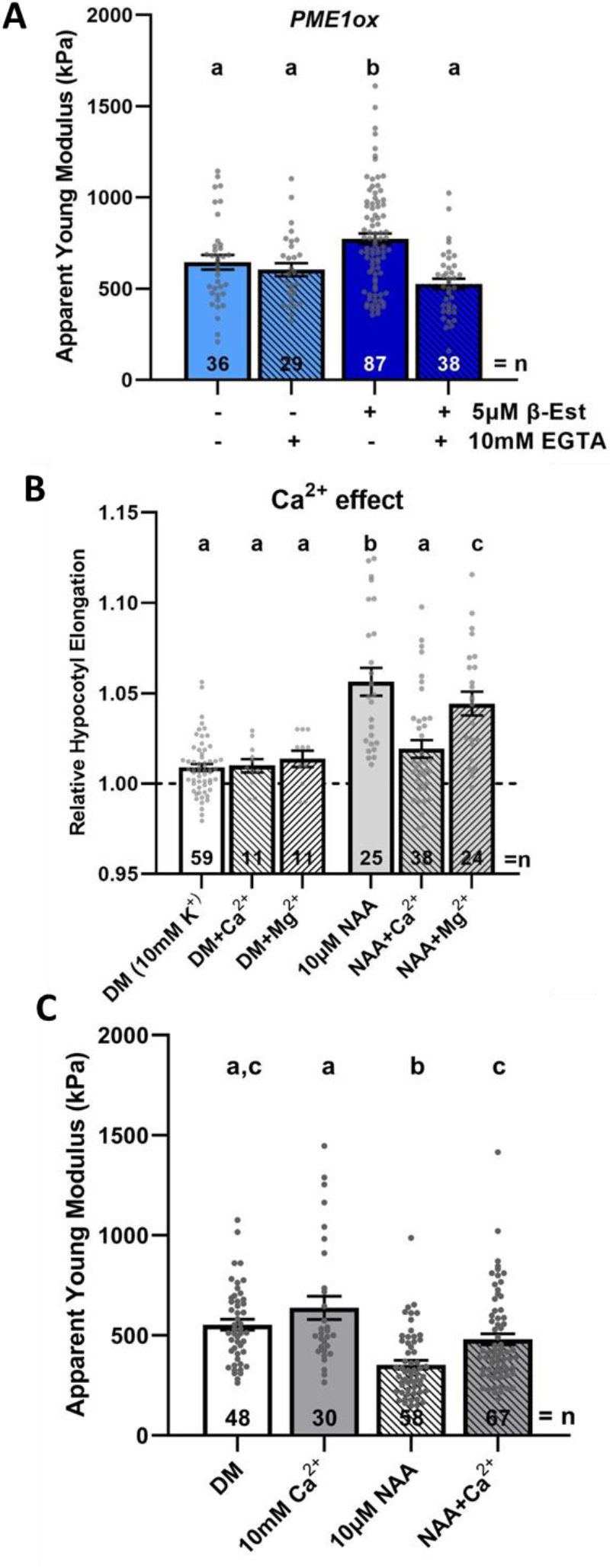
Calcium interferes with auxin-triggered hypocotyl elongation and CW softening. A. Average *E_a_* quantification of longitudinal CWs of hypocotyl epidermis in 3 day-old hypocotyl segments from *PME1ox* line non- or treated for 24 h with β-EST and incubated or not for 2 h in medium supplemented with calcium chelator (ethyleneglycoltetraacetic acid, EGTA). B. Relative hypocotyl segment length after 3 h of indicated treatments. DM was modified adding 5mM CaCl_2_ or MgCl_2_ (together with 5mM KCl) as ion interfering treatments. C. Average *E_a_* quantified on longitudinal epidermal CWs of 3 day-old hypocotyl segments incubated for 2 h in depletion medium supplemented with auxin (1-naphthaleneacetic acid, NAA) and Ca^2+^ as indicated. Dashed line indicate relative to initial length of hypocotyl segments. Bars represent average ±*SE*, dots represents individual data points. N indicates the total number of scans performed and analyzed (A and B) and the hypocotyls segments quantified (C).Tukey-Kramer test was performed and significant differences are shown in the letters above each bar.

**Figure 5.**
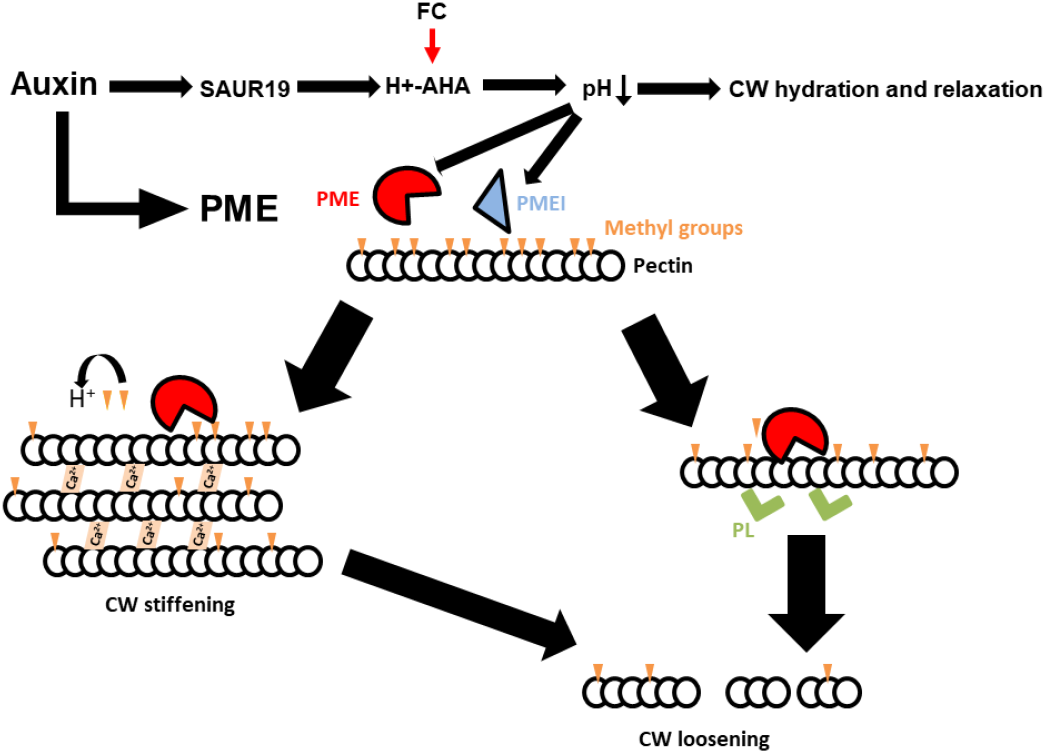
Model for dual role of PME activity. Hormonal treatments converge on acidification of apoplast, which trigger a CW relaxation. In parallel, short-term PME activation might generate pectin de-methylesterification pattern that enhances calcium-mediated cross-linking and stiffening of CW, whereas long-lasting PME activity might induce pectin degradation and CW softening. Partially adapted from Hocq et al., 2017b.

Using maize coleoptiles it has been shown that Ca^2+^ added to the medium interferes with auxin-induced elongation (Tode and Lüthen, 2001). Consistently, we observed that Ca^2+^ added into the depletion medium attenuated auxin triggered elongation of Arabidopsis hypocotyl segments. When compared to Ca^2+^, divalent Mg^2+^ ions interfered with auxin-induced elongation significantly less (Fig. 4B). However, whether Ca^2+^ inhibition of auxin-induced hypocotyl elongation involves alteration of CW mechanical properties has not been investigated. AFM measurements of hypocotyl segments revealed significantly increased CW stiffness in epidermal cells of hypocotyls incubated in the medium supplemented with auxin in the presence of Ca^2+^, when compared to control hypocotyls exposed to auxin only (Fig. 4C), suggesting that Ca^2+^ availability might significantly affect auxin-mediated softening of CWs.

Altogether, our data show that levels of extracellular calcium play a decisive role in auxin-mediated softening of the CWs and hypocotyl elongation. Furthermore, they support a model in which a short-term PME activity generates a de-methylation pattern of HGs, which is prone to cross-linking by Ca^2+^ that promotes stiffening of CWs.

## CONCLUSIONS

The relation between CW composition and its mechanical properties is still a big unsolved enigma. CW remodeling is a dynamic process that is an integral part of regulatory mechanisms controlling growth and development of plant organs. Modulation of CW biophysical properties is a result of a complex regulatory network encompassing multiple plant hormones, CW remodeling enzymes, calcium ions and pH (Wolf et al., 2012; Hocq et al., 2017b).

Here, we show that acidification is a necessary step for CW relaxation in hypocotyl cells. When acidification by hormones is compromised by an alkaline environment, softening of CWs and growth is significantly attenuated, indicating that it is a shared mechanism among several plant hormones that promote hypocotyl elongation.

Pectin is an important component of CW. A recent report using 3D-STORM microscopy applied on cotyledon pavement cell suggested that degree of methylation of HGs might determine quaternary structure that defines intrinsic pectin swelling properties. These observations were summarized in the “expanding beam model” proposing that local HG de-methylesterification leads to nanofilament radial swelling, which is caused by conversion between quaternary structures with different packaging (Haas et al., 2020, 2021). The author showed that de-methylesterification of HG alone is sufficient to induce expansion in cotyledon pavement cells. In such context, the balanced activities of PME and PMEI enzymes should be a key factors regulating cell growth. Accordingly, previous reports have demonstrated that PME and PMEI control CW biophysical properties (Bou-Daher et al., 2018; Peaucelle et al., 2015; Jonsson et al., 2021), suggesting that demethylesterification of HGs is an essential determiner of cell expansion and tissue growth.

However, experimental observations showing that enhanced PME activity might lead to both, reduced or increased CW stiffness, raised a question about underlying mechanisms. This led to formulation of a model proposing that the distinct outputs might be dependent on the pattern of pectin-de-methylation by PME (Hocq et al., 2017b). Our experiments corroborate the model, in which PME long-term or high activity induces softening of CWs most probably because loss of methyl groups allows pectin lyases and other degradation enzymes to break pectin fibers, whilst short-term or moderate PME activity lead to CW stiffening as result of pectin cross-linking by calcium.

Notably, hypocotyl elongation assays are typically performed under calcium depletion, which might create artificial conditions under which pectin is not able to form new calcium cross-links due deficiency of calcium ions. Indeed, our results show that adding calcium might interfere with auxin-triggered CW softening and elongation processes. Furthermore, it is proposed that acidification and CW relaxation is correlated with calcium import (Cho et al., 2012; Conn et al., 2011) as result of cell expansion that might stretch the plasma membrane and open stretch-activated calcium channels.

Overall, we have demonstrated that acidification and growth processes mediated by several hormones converge on the CW softening process and that regulation of the PME activity tightly linked with calcium levels are essential factors determining CW mechanical properties. Further studies characterizing the effects of expansin proteins on the CW composition and mechanical properties might shed light on the complex network of CW remodeling in growth processes.

## Material and Methods

### Plant material

*Arabidopsis thaliana* plants were grown in a growth chamber at 21°C under white light (W), which was provided by blue and red LEDs (70-100 μmol m^-2^s^-1^ of photosynthetically active radiation). The transgenic lines have been described elsewhere: *HS::axr3-1* (Knox et al., 2003), *35S::GFP-SAUR19* (Spartz et al., 2012), R2D2 (Liao et al., 2015), ApopH (Fendrych et al., 2016). The *HS::axr3-1* plants were heat-shocked at 37°C for 40 min in aluminum- wrapped petri dishes; experiments were started 1.5hr after the end of the heat shock.

All cloning and transformation procedure was conducted using Gateway™ (Invitrogen) technology, with the sequences of all used vectors available online (https://gateway.psb.ugent.be/), similarly as described before (Hurný et al., 2020). Inducible *PMEox* lines were generated by PCR amplification of cDNA from 3 days-old seedlings and insertion on pDONR221 ™ B1-B2 Gateway vector. Primers used for the cloning can be found in Table S1. The inserts were confirmed by PCR and sequencing. Afterwards, the clones were recombined into pDEST™ R4-R3 pB-rPPT, together with the promoter from pUBQ10-XVE-P4P1 and the tag from pEN-R2-3xHA-L3 vectors. All transgenic plants were generated by the floral dip method in Columbia (Col-0) background and transformants were selected on plates with respective antibiotic. Induction by β-Est was done by transferring seedlings in AM plate with indicated concentrations of β-Est at indicated times.

### Growth conditions

Seeds were sterilized in 5% bleach for 10 min and rinsed with sterile water before plating on half-strength Murashige and Skoog (MS) medium (Duchefa) with 1% sucrose, 1% agar (pH 5.7). Seeds were stratified for 3-4 days at 4°C, exposed to light for 2-4 hours at 21°C, and cultivated in the growth chamber under appropriated light conditions at 21°C (wrapped in aluminum foil and disposed in a cardboard box for dark growth, or into light in the same chamber).

### Elongation assays

Hypocotyl elongation assays were performed as described elsewhere (Fendrych et al., 2016) with minor modifications. Hypocotyl segments were obtained from etiolated seedlings ~72h-old (starting after transferring plates into growth-room), by removing the hook and the lower half part of the hypocotyl (and root) using a razor blade in a darkness (a green LED light was used for illumination). The hypocotyl segments were placed on the surface of the Depletion Medium (DM, 10 mM KCl, 1 mM MES, pH 6,0 using KOH, 1.5% phytagel) and kept in darkness for at least 20 min. pH was adjusted using HCl or NaOH 10mM. Calcium and magnesium were added as CaCl_2_ or MgCl_2_ (Sigma-Aldrich) at indicated concentrations. Then the segments were transferred to a new plate with depletion medium and the supplemented hormones or inhibitors as specified in the figure legends. The samples were placed on a flatbed scanner (Epson perfection V370) and imaged through the layer of phytagel, a black filter paper was placed above the dishes to improve the contrast of the images. Samples were scanned immediately after transfer (used as initial length) and, unless other specification, after 3 h of incubation in the respective treatment.

### Root lengths

ImageJ software (http://rsb.info.nih.gov/) was used on digital images to measure the root length of 3 days-old dark grown seedlings non- or treated with 5 μM β-Est for 48 h (24 h old seedlings were transferred to a new AM plate containing or not β-Est and incubated for 48 h in darkness).

### Chemicals used

Auxin, 1-naphthaleneacetic acid (NAA), epibrassinolide (eBL), Gibberellic acid (GA), Fusicoccin (FC) and β-Estradiol (β-Est) and ethylene glycol-bis(β-aminoethyl ether)-N,N,N’,N’-tetraacetic acid (EGTA) were ordered from Sigma-Aldrich. NAA and eBL were used at 10 mM, GA was dissolved to 100mM stock concentration and FC at 1 mM in EtOH. β-Est was dissolved in water at 10 mM. EGTA was prepared in stock solution at 0.5 M in water.

### Confocal imaging

Confocal laser-scanning micrographs were obtained with a Zeiss LSM800 with a 488-nm argon laser line for excitation of GFP fluorescence. Emissions were detected between 505 and 580 nm. Using a 20x air objective, confocal scans were performed with the pinhole at 1 Airy unit. Localization was examined by confocal z-sectioning. Each image represents either a single focal plane or a projection of individual images taken as a z-series. Z-stacking was performed collecting images through cortex and epidermal layers. Full z-stack confocal images were 3D-projected using ImageJ software. At least five seedlings were analyzed per treatment. Images were processed and quantified in ImageJ. For ApopH regions of interest (ROIs) were defined at plasma membrane using manual line of ImageJ tool and for R2D2 individual nuclei were selected as ROIs using manual selection round shape.

### AFM measurements and Apparent Young’s Modulus Calculations

The AFM data were collected and analyzed as described elsewhere with minor changes (Peaucelle et al., 2015; Hurný et al., 2020; Velasquez et al., 2021). To examine extracellular matrix properties we suppressed turgor pressure by immersion of the seedlings in a hypertonic solution (10% mannitol) for at least 20min before examination. Three day-old seedlings grown in darkness (in normal AM plate or indicated treatment) or hypocotyl segments treated as described for elongation assays were placed in microscopy slides and immobilized using double-glued side tape. We focused on the longitudinal CWs (parallel to growth axis, but perpendicular to the organ surface), and its extracellular matrix. To ensure proper indentations, especially on the regions in the bottom of the doom shape between two adjacent cells, we used cantilevers with long pyramidal tip (14-16 μm of pyramidal height, AppNano ACST-10), with a spring constant of 7.8 N/m. All cantilevers were calibrated prior to biomechanical experiments. The stiffness of the cantilever was calibrated by thermal noise measurement and the sensitivity was calibrated by force-distance curve measurement with a microscopy glass slide (setpoint value 1.5 V). The instrument used was a JPK Nano-Wizard 4.0 and indentations were kept to <10% of cell height (typically indentations of 100-200 nm depth and 500 nN force). Typically, three scan-maps per sample were taken over an intermediate region of the hypocotyl, using a square area of 25 x 25 μm, with 16 x 16 force-indentation measurements. The lateral deflection of the cantilever was monitored and in case of any abnormal increase the entire data set was not used for analysis. The apparent Young’s modulus (*E_a_*) for each force-indentation experiment was calculated using the approach curve (to avoid any adhesion interference) with the JPK Data Processing software (JPK Instruments AG, Germany), based on Herz model adjusted to pyramidal tip geometry. To calculate the average *E_a_* for each longitudinal wall, the *E_a_* was measured over the total length of the extracellular region using masks with Gwyddion 2.45 software (at least 20 points were taken in account). The pixels corresponding to the extracellular matrix were chosen based on topography and Young Modulus maps. For topographical reconstructions, the height of each point was determined by the point-of-contact from the force-indentation curve. A standard t-test was applied to test for differences between genotypes/treatments. Total number of scans performed and analyzed is indicated in the figures (indicated as n).

### Expression analysis by RT-qPCR

Around 60 seedlings (72h old dark grown of indicated lines, transferred after 48 h to AM plates containing or not 2 μM β-Est) were harvested and frozen in liquid nitrogen. Tissue was ground using a ball mill (model MM400; Retsch) with 4-mm diameter balls in a 2-ml Eppendorf (Hamburg, Germany). Total RNA was isolated using Monarch kit (New England Biolabs) according to manufacturer protocol (DNase treatment was included during extraction protocol). cDNA was prepared from 1 μg of total RNA with the iScript cDNA Synthesis Kit (Biorad), diluted 10 times and 1 μl was used in 5-μl PCR reaction on a LightCycler 480 (Roche Diagnostics) with Luna Master Mix (New England Biolabs) according to the manufacturer’s instructions. As control, non-RT-treated samples were included to test the purity of the cDNA. All experiments were done with three technical replicates and three biological samples. The PP2A gene (At1g69960) was used as a control for normalizations. Primer sequences can be found in Table S1.

### Western-blot analysis

Around 60 seedlings (72h old dark grown of indicated lines, transferred after 48 h to AM plates containing or not 2 μM β-Est) were harvested and frozen in liquid nitrogen. Frozen tissue was ground with stainless steel 4-mm diameter balls in a 2-ml Eppendorf (Hamburg, Germany) tube using a ball mill (model MM400; Retsch). Extraction buffer [100 mM Tris–HCl (pH 7.5), 25% (w/w, 0.81 M) sucrose, 5% (v/v) glycerol, 10 mM ethylenediaminetetraacetic acid (EDTA, pH 8.0), 10 mM ethyleneglycoltetraacetic acid (EGTA pH 8.0), 5 mM KCl, and 1 mM 1,4-dithiothreitol (DTT); Abas and Luschnig, 2010] was added to the frozen tissue. After Bradford quantification, 20 μg of protein were diluted in 20μl of buffer and prepared for electrophoresis by adding 5μl of loading buffer (5xSDS) and incubating at 45°C for 5min. Run was performed in a commercial 10% Mini-PROTEAN^®^ TGX™ Precast Protein Gels (Bio-Rad) at 35mA. Transference was done with semidry system with a Trans-blot turbo transfer pack PVDF (Bio-Rad). The blot was washed in TBST buffer with 5% Milk Powder and 0.05% Tween20, and blocked over/night in the same buffer at 4°C. Hybridization was done with anti-HA-HRP antibody (monoclonal antibody from Sigma, dilution 1:7000) in TBST for 2h at room temperature. The blot was washed three times in TBST and one in water, previous the visualization by SuperSignal West Femto Maximum Sensitivity Substrate kit (Thermo Scientific) and exposure to Amersham Imager 600 (GE Healthcare).

## Accession Numbers

Sequence data from this article can be found in the Arabidopsis Genome Initiative or GenBank/EMBL databases under the following accession numbers: At1g53840 (*PME1*), At5g47500 (*PME5*).

## Acknowledgements

We acknowledge Jaume F. Martínez García for *phyAphyB* mutant seeds. This work was supported by grants from the European Research Council (Starting Independent Research Grant ERC-2007-Stg-207362-HCPO to E.B.) and M.G. was recipient of an IST Interdisciplinary project (IC1022IPC03). We acknowledge CF Nanobiotechnology of CIISB, Instruct-CZ Centre, supported by MEYS CR (LM2018127).

## Author contributions

Conceived and designed the experiments: MG, EH and EB. Performed the experiments: MG; JMCL contributed with western-blot and immuno assays and discussion of whole manuscript; NZ performed western-blot and elongations assays; JP and PS contributed in AFM experiments. Analyzed the data: MG, EH and EB. Wrote the manuscript: MG and EB with inputs from all authors.

## Competing interests statement

The authors declare no competing financial interests.

## Notes

### Competing Interest Statement

The authors have declared no competing interest.

### Summary of Updates

Shortened and simplified version of the manuscript, merging Results&Discussion, figures and text adjusted and corrected.

