## Supplemental Figures and Table S1 for "Dual role of Pectin Methyl Esterase activity in the regulation of plant cell wall biophysical properties"

### Supplementary Figure 1

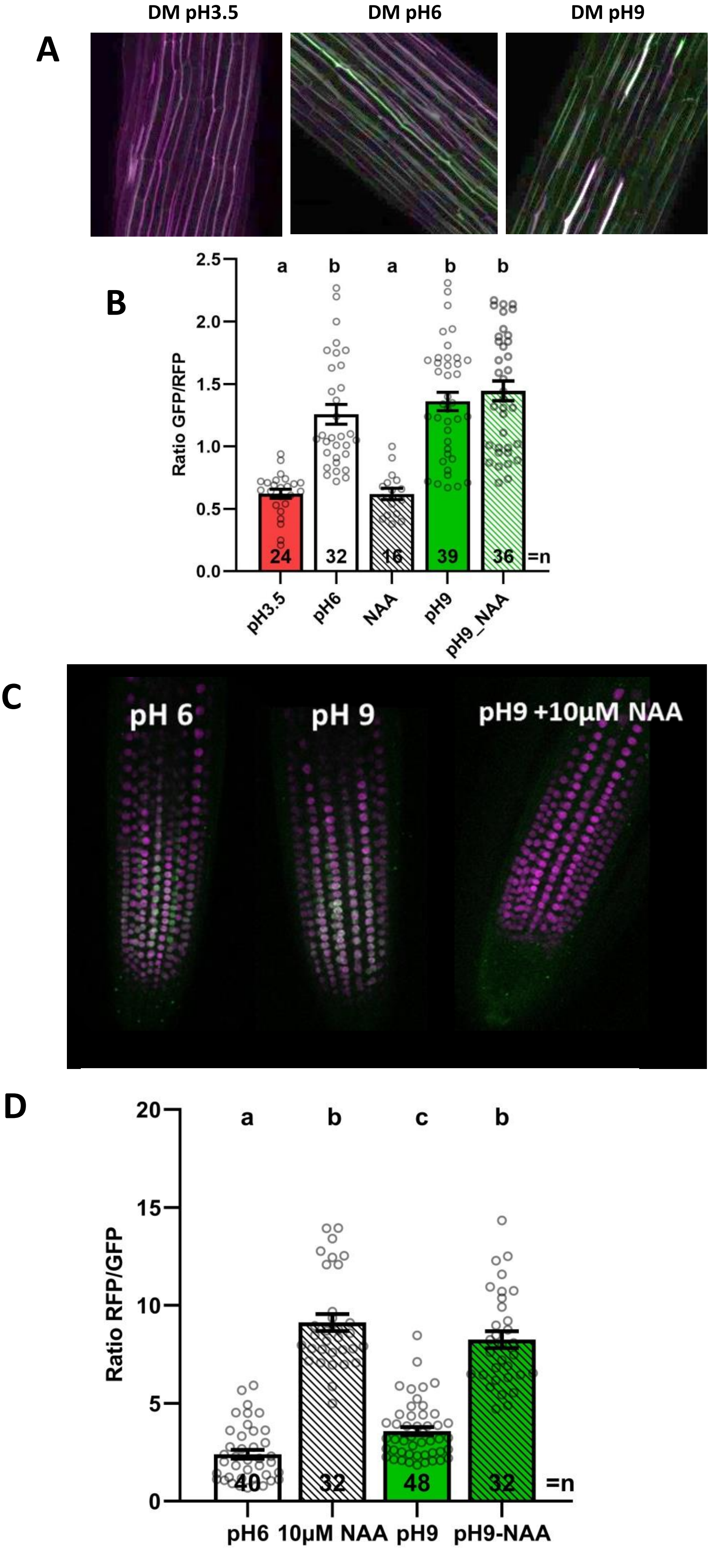

**Supplementary Figure 1. Effect of pH buffered media on apoplastic region and auxin signaling.** A and B. Hypocotyl segments of 3 days-old seedlings expressing ApopH marker line (GFP/RFP) incubated in buffered depletion medium (DM) with pH as indicated for 2 h. Maximum intensity Z-stack projection images (A) and apoplast pH quantified as GFP/RFP intensity ratio (B). C and D. 3 days-old roots expressing the *R2D2* auxin signaling reporter 2 h after indicated treatment. Maximum intensity Z-stack projection images of root meristematic zone (C) and quantification of auxin response as RFP/GFP signal ratio (D). Bars represent average ±*SE*, dots represents individual data points. N indicates the total number of cells quantified. Tukey-Kramer test was performed and significant differences are shown as the letters above each bar.

### Supplementary Figure 2

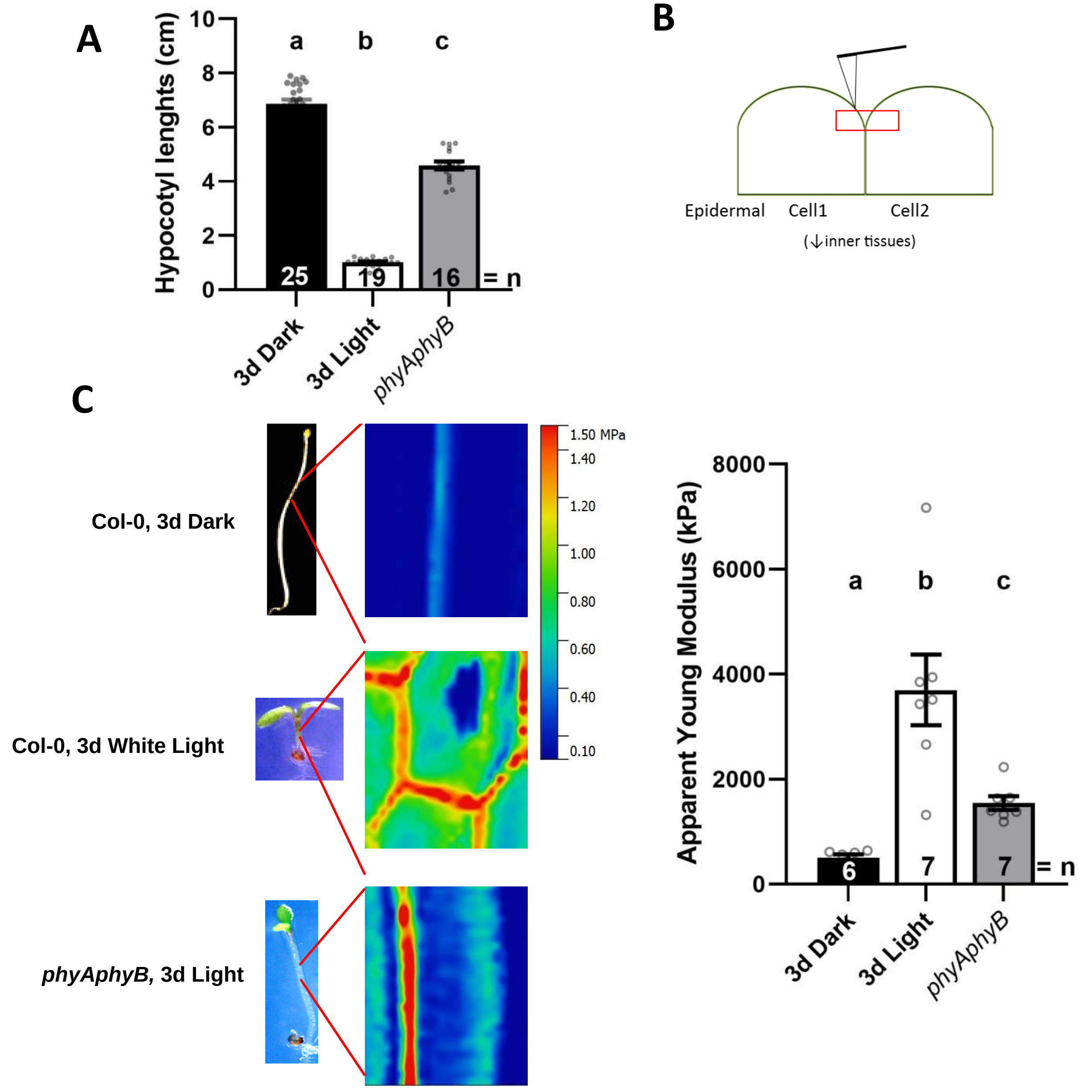

**Supplementary Figure 2. Atomic force microscopy (AFM) analyses of cell wall stiffness in hypocotyl epidermal cells .** A. Hypocotyl lengths of 3 days-old wild-type (Col-0) and *phyAphyB* seedlings grown in light or darkness. B. Scheme of the experimental setup used for AFM measurements. Longitudinal cell walls between two hypocotyl epidermal cells, parallel to growth axis (indicated by red square), were scanned. C. Measurements of Average Apparent Young ($E_a$ ) (right) and corresponding heat-maps (middle) of epidermal cells in hypocotyls of Col-0 and *phyA,phyB* (left) . Scale bar = 5µm. Bars represent average ±*SE*, dots represents individual data points. N indicates the total number of hypocotyls measured (A) or individual longitudinal cell wall scans performed and quantified (C). Tukey -Kramer test was performed and significant differences are shown as the letters above each bar.

### Supplementary Figure 3

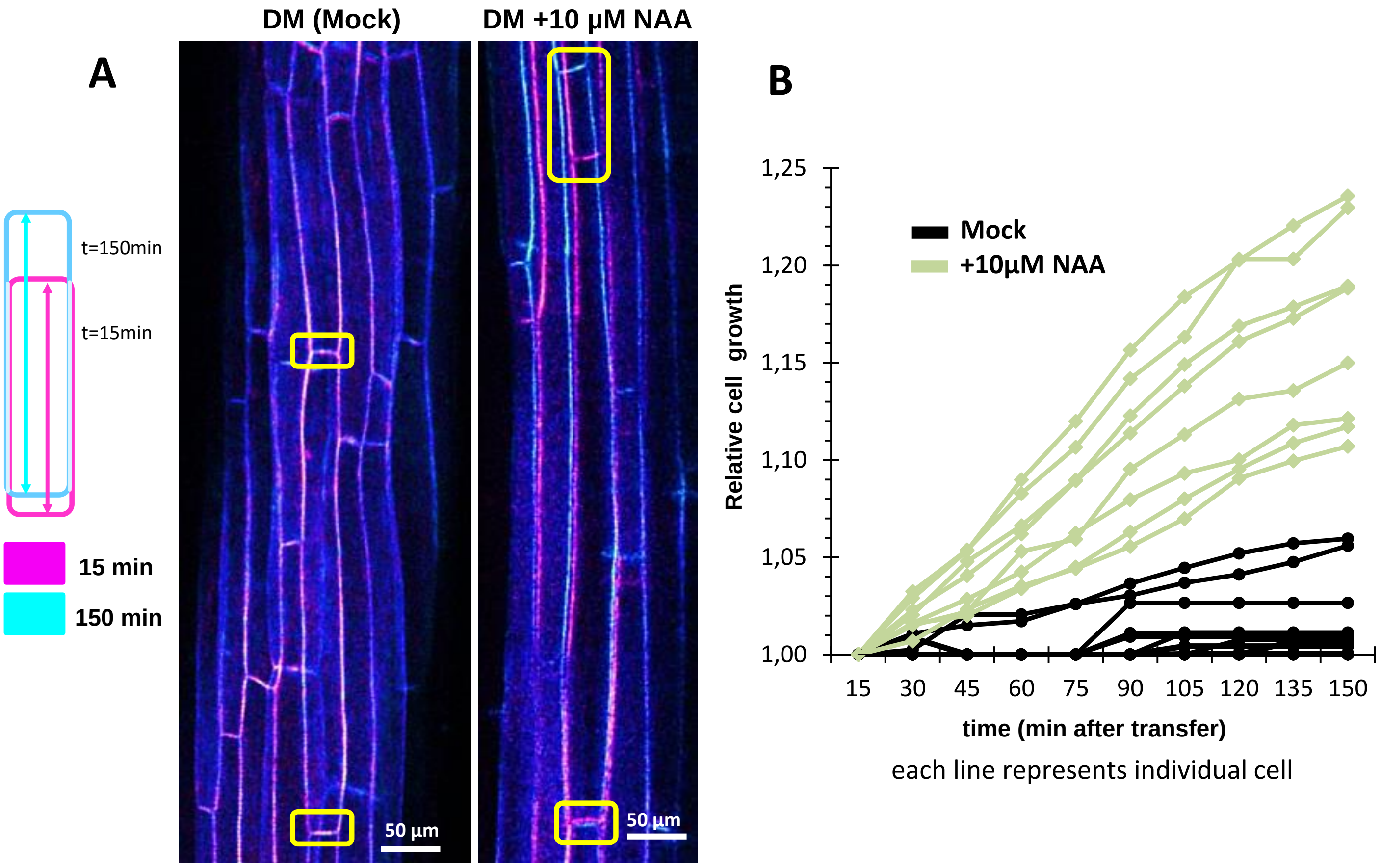

**Supplementary Figure 3. Auxin promotes hypocotyl epidermal cell elongation.** A. A color code representation of hypocotyl epidermal cell elongation. Cell length at 15 min and 150 min, magenta and cyan, respectively, after incubation in the depletion medium (DM) and DM supplemented with auxin (1-naphthaleneacetic acid, NAA). Images are maximum intensity Z-stack projection of 3 day-old hypocotyl segments from seedlings expressing *35S::PIP2-GFP* plasma membrane reporter. Yellow squares remark areas were cell elongation can be observed. Scale bar = 50 µm. B. Time course of epidermal cell length increase. Epidermal cells in the middle part of hypocotyl segments were monitored and measured in indicated time intervals (cells from 3 different hypocotyls are shown).

### Supplementary Figure 4

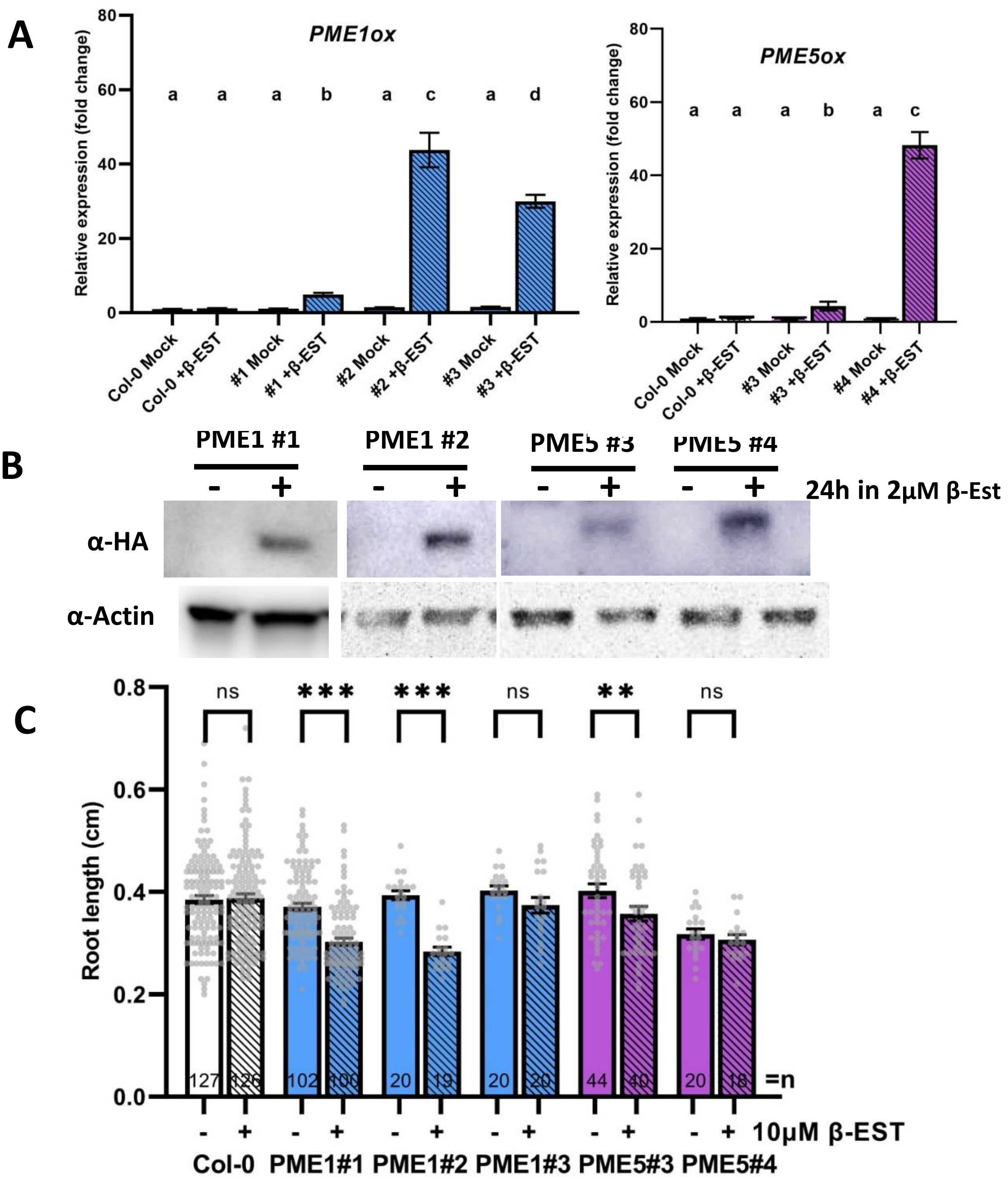

**Supplementary Figure 4. Expression and phenotype analyses of *PMEoxs* lines.** A. Relative expression levels of *PME1ox* and *PME5ox* analysed by RT-qPCR in several independent homozygous transgenic lines. Expression of *PME*1 and *PME5* induced by treatment with 2 µM β-Est for 24 h. Expression monitored in whole 3 days-old dark grown seedlings. Tukey-Kramer test was performed and significant differences are shown as the letters above each bar. B. PME1-HA and PMA5-HA protein detected by Western-Blot analysis in several independent *PMEox*s transgenic lines induced by same treatments as mentioned before. α-actin monitored as a protein loading control. C. Root lengths several independent *PMEox*s transgenic lines. 3 days-old dark grown seedlings non- or treated with 5 µM β-Est for 48 h. Bars represent average *±SE*, dots represents individual data points. N indicates the total number of roots quantified. Direct pair significant differences are indicated as **P < 0.01 and ***P < 0.001 (t-test).

### **Supplementary Table S1.** List of primers used.

| Name | Sequence | Gene\Seq | Purpuse |
|---|---|---|---|
| **MGR108_PP2A F** | TAACGTGGCCAAAATGATGC | AT1G69960 | qPCR HK gene |
| **MGR109_PP2A R** | GTTCTCCACAACCGCTTGGT | AT1G69960 | qPCR HK gene |
| **MGR112_PME1F** | ACGGTGATTGGAGCAGTGG | AT1G53840 | qPCR PME1 |
| **MGR113_PME1R** | CTGCGTTACATCTGAACCCG | AT1G53840 | qPCR PME1 |
| **MGR114_PME5F** | TCACCGTCTCACTTAACGGC | AT5G47500 | qPCR PME5 |
| **MGR115_PME5R** | TAGCGGTCACATCCCTACCA | AT5G47500 | qPCR PME5 |
| **MGR90_PME1Fw** | GGGGACAAGTTTGTACAAAAAAGCAGGCTCC_ATGGATTCAGTGAACTCCTTCAAAGG | AT1G53840 | Cloning attB1 PME1 |
| **MGR91b_PME1Rv** | GGGGACCACTTTGTACAAGAAAGCTGGGTGTA_AGATAGCTGATTGATCACTCCTGTTGC | AT1G53840 | Cloning attB2 PME1 |
| **MGR92_PME5Fw** | GGGGACAAGTTTGTACAAAAAAGCAGGCTCC_ATGGCGCAACTTACTAATTCCCTC | AT5G47500 | Cloning attB1 PME5 |
| **MGR93b_PME5Rv** | GGGGACCACTTTGTACAAGAAAGCTGGGTGTA_AGCATCTCGAGGAGCGATCC | AT5G47500 | Cloning attB2 PME5 |
| **MGR111_HARv** | CCGCATAGTCAGGAACATCG | HA tag | Genotypings |
